# Skeletal muscle DNA methylation and mRNA responses to a bout of higher versus lower load resistance exercise in previously trained men

**DOI:** 10.1101/2022.11.01.514721

**Authors:** Casey L. Sexton, Joshua S. Godwin, Mason C. McIntosh, Bradley A. Ruple, Shelby C. Osburn, Blake R. Hollingsworth, Nicholas J. Kontos, Philip J. Agostinelli, Andreas N. Kavazis, Tim N. Ziegenfuss, Hector L. Lopez, Ryan Smith, Kaelin C. Young, Varun B. Dwaraka, Andrew D. Fruge, Christopher B. Mobley, Adam P. Sharples, Michael D. Roberts

## Abstract

We sought to determine the skeletal muscle genome-wide DNA methylation and mRNA responses to one bout of lower-load (LL) versus higher-load (HL) resistance exercise. Trained college-aged males (n=11, 23±4 years old, 4±3 years self-reported training) performed LL or HL bouts to failure separated by one week. The HL bout (i.e., 80 Fail) consisted of four sets of back squats and four sets of leg extensions to failure using 80% of participants estimated one-repetition maximum (i.e., est. 1-RM). The LL bout (i.e., 30 Fail) implemented the same paradigm with 30% of est. 1-RM. Vastus lateralis muscle biopsies were collected before, 3 hours, and 6 hours after each bout. Muscle DNA and RNA were batch-isolated and analyzed using the 850k Illumina MethylationEPIC array and Clariom S mRNA microarray, respectively. Performed repetitions were significantly greater during the 30 Fail versus 80 Fail (p<0.001), although total training volume (sets x reps x load) was not significantly different between bouts (p=0.571). Regardless of bout, more CpG site methylation changes were observed at 3-versus 6-hours post exercise (239,951 versus 12,419, respectively; p<0.01), and nuclear global ten-eleven translocation (TET) activity, but not global DNA methyltransferase activity, increased 3- and 6-hours following exercise regardless of bout. The percentage of genes significantly altered at the mRNA level that demonstrated opposite DNA methylation patterns was greater 3- versus 6-hours following exercise (~75% versus ~15%, respectively). Moreover, high percentages of genes that were up- or downregulated 6 hours following exercise also demonstrated significantly inversed DNA methylation patterns across one or more CpG sites 3 hours following exercise (65% and 82%, respectively). While 30 Fail decreased DNA methylation across various promoter regions versus 80 Fail, transcriptome-wide mRNA and bioinformatics indicated that gene expression signatures were largely similar between bouts. Bioinformatics overlay of DNA methylation and mRNA expression data indicated that genes related to “Focal adhesion”, “MAPK signaling”, and “PI3K-Akt signaling” were significantly affected at the 3- and 6-hour time points, and again this was regardless of bout. In conclusion, extensive molecular profiling suggests that post-exercise alterations in the skeletal muscle DNA methylome and mRNA transcriptome elicited by LL and HL training bouts to failure are largely similar, and this could be related to equal volumes performed between bouts.

## INTRODUCTION

Resistance training increases myofiber hypertrophy, whole-tissue hypertrophy, and strength [1]. Training with higher loads (e.g., ~80%+ of a person’s one repetition maximum, or 1RM) and lower volumes (e.g., ~8-12 repetitions per set) generally increases force production capabilities versus lower load training with higher volumes (e.g., 20+ repetitions per set with 30-60% 1RM) [2–5], although equivocal evidence exists [6,7]. Notwithstanding, most studies to date support that a wide range of loads ≥30 1RM can promote a similar magnitude of skeletal muscle hypertrophy if training is performed to failure [5–13].

Although the effects of volume-load manipulations on strength and hypertrophy have been extensively investigated over recent years, much less attention has been given to the potential divergent molecular responses that may occur between higher load and lower load training. Initial findings by Mitchell et al. [5] and Haun et al. [14] suggested that one bout of lower-load and higher-load failure training similarly elevated post-exercise anabolic signaling events in skeletal muscle (i.e., mTORC1 phosphorylation markers). However, more recent research provides evidence supporting that different muscle-molecular adaptations may occur between training paradigms. For instance, Lim and colleagues [10] reported that 10 weeks of 30% 1RM training to failure affected markers associated with mitochondrial biogenesis and remodelling versus 80% 1RM training to failure. Haun et al. [15] reported that a six-week lower load and higher volume resistance training program in previously trained men elicited a significant upregulation in sarcoplasmic proteins associated with glycolysis measured via proteomics. In a separate cohort of previously trained men, Vann et al. [16] reported that a 10-week higher load and moderate-to-low volume training program did not alter the sarcoplasmic proteome or promote the glycolytic protein adaptations observed by Haun and colleagues. Vann et al. [4] subsequently examined how six weeks of unilateral lower load versus higher load unilateral leg resistance training affected select molecular outcomes in a third cohort of previously trained men. While shotgun proteomics were not performed, these authors reported that six-week integrated non-myofibrillar protein synthesis rates were significantly greater in the lower load versus higher load trained leg. This collective evidence has led our laboratory to hypothesize that lower load and higher volume resistance training promotes metabolic adaptations relative to higher load training [17].

Unique molecular signaling events induced by exercise training promote alterations in mRNA expression [18], and several lines of evidence suggest that epigenetic modifications to DNA is a mechanism involved in this process [19–21]. DNA methylation is one of the most studied epigenetic mechanisms in humans, and recent research supports that a variety of exercise modes, including resistance training, can alter DNA methylation status across various gene promoters [22–24]. In mammalian species, DNA methylation primarily occurs at cytosine and guanine dinucleotide-rich sites (CpGs). DNA methylation is catalyzed by DNA methyltransferase (DNMT) enzymes, and DNA de-methylation is catalyzed by ten-eleven translocation methylcytosine dioxygenases (TETs) [20,25]. Increased metabolite flux through glycolysis and the citric acid cycle are thought to alter TET activity [26], and affect the pool of methyl groups available for donation during the methylation process [25]. Current dogma indicates that in the majority of cases decreased methylation (hypomethylation) of CpG sites, particularly if occurring in promoter regions, contributes to increased gene expression whereas increased methylation (hypermethylation) contributes to gene silencing [19,21].

It is plausible that divergent genome-wide DNA methylation and mRNA expression signatures induced by higher and lower load resistance training could contribute to some of the unique muscle-molecular phenotypes that have been previously observed between these paradigms. However, neither phenomenon have been previously investigated. Therefore, the purpose of this study was to examine how an acute bout of higher-versus lower load resistance exercise to failure affected genome-wide DNA methylation and mRNA transcription profiles in skeletal muscle. Moreover, we sought to determine whether DNA methylation events after each mode of training overlapped with mRNA responses. Herein, lower-load and higher-load resistance exercise to failure utilized loads that were 30% 1-RM (30 Fail) and 80% 1-RM (80 Fail), respectively. Given previous findings, we hypothesized that 30 Fail training would elicit greater DNA hypomethylation relative to 80 Fail training, and this would correspond to an increased expression of mRNAs associated with mitochondrial biogenesis, mitochondrial remodelling, and glycolysis.

## METHODS

### Ethical Approval and Pre-screening

This study was conducted with prior review and approval from the Auburn Institutional Review Board (IRB approval #: 20-081 MR 2003), and in accordance with the most recent revisions of the Declaration of Helsinki except not being pre-registered as a clinical trial. The participants recruited for this study were males from the local community that met the following criteria: i) 18-35 years old, ii) body mass index ([BMI] body mass in kilograms/ height in meters^2^) not exceeding 35 kg/m^2^, iii) no known cardio-metabolic disease (e.g., obesity, diabetes, hypertension, heart disease) or any condition contraindicating participation in resistance training or donating muscle biopsies, iv) have participated in lower-body training at least once per week over the last six months, v) have a self-reported barbell back-squat one repetition maximum of ≥ 1.5x body weight. Following verbal and written consent, participants proceeded with study procedures described below.

### Study Design

Figure 1 provides a schematic of the study design. Participants completed a within-subject design that included a total of five visits to the laboratory. At Visit 1, an informed consent and health history questionnaire were completed. This was followed by barbell back squat and knee extension strength tests as well as body composition testing for phenotyping purposes. At Visit 2 (~7 days following Visit 1), participants completed a battery of baseline (PRE) tests that included measures of height, weight, and vastus lateralis (VL) thickness. Participants also donated a muscle biopsy from the VL prior to the completion of a single bout of resistance exercise using a randomly assigned experimental load (i.e., 30% or 80% estimated 1RM). At both 3 hours (3h POST) and 6 hours (6h POST) following the exercise bout, participants donated additional muscle biopsies from the VL. Seven days following Visit 2, participants completed another bout of resistance exercise using the experimental load not completed at Visit 2 (i.e., if Visit 2 was lifting to failure at 30% 1RM, then Visit 4 implemented 80% 1RM loads). As with Visit 2, VL biopsies were obtained at PRE, 3h POST, and 6h POST. Visits 3 and 5 were follow-up visits to ensure biopsy sites were properly healing. The paragraphs below provide more expanded details on testing procedures.

**Figure 1.**
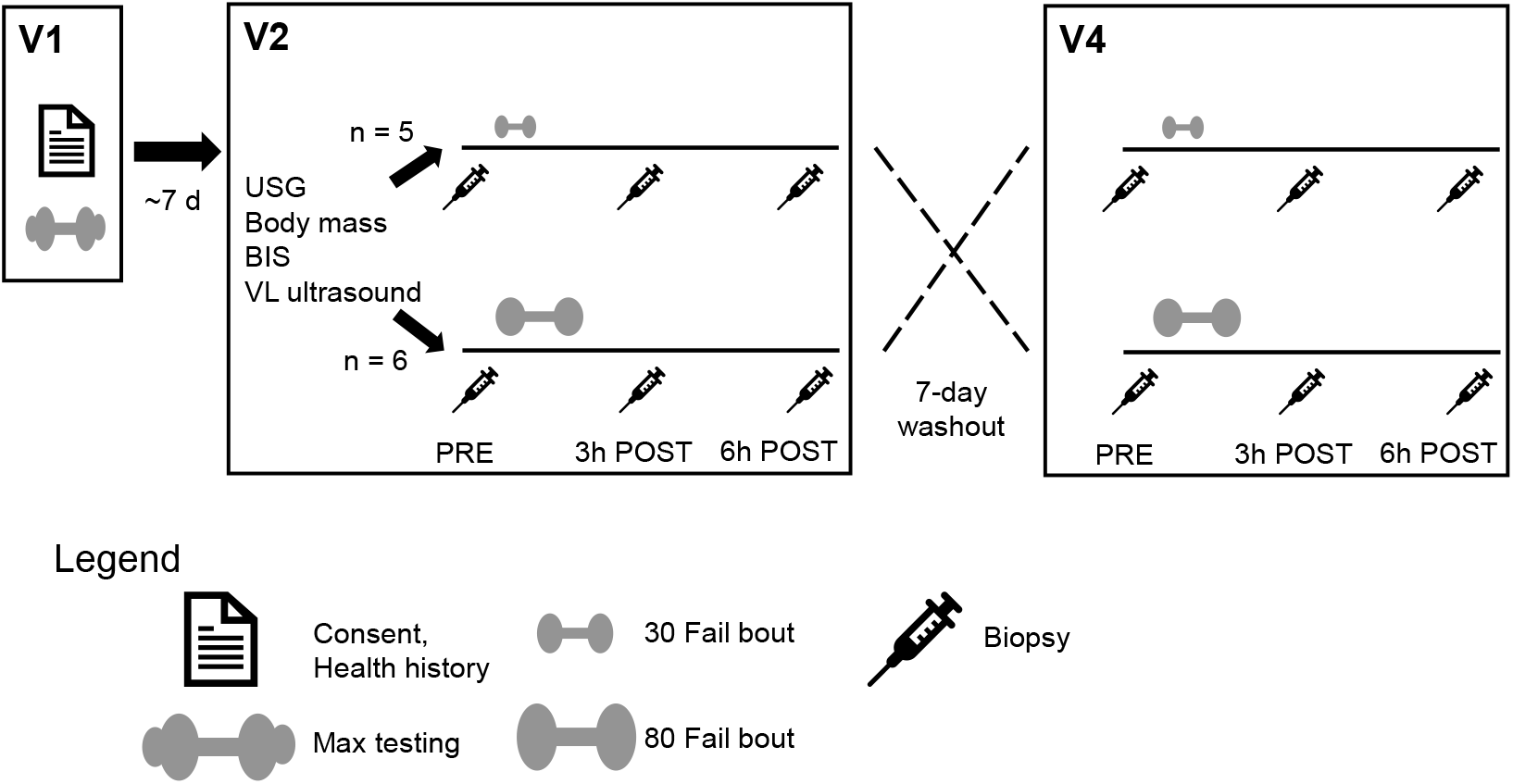
Study design **Legend:** This schematic represents the timeline for participant consent, max testing, PRE-phenotype testing, experimental resistance exercise bouts, cross-over washout period, and timepoints for all biopsies. Abbreviations: 30 Fail, 30% one repetition maximum to failure; 80 Fail; 80% one repetition maximum; BIS, bioelectrical impedance spectroscopy (for body composition); USG, urine-specific gravity testing; V1/V2/V4, Visits 1/2/4; VL, vastus lateralis.

### Testing Sessions

#### Estimated 1-RM testing

During visit 1, participants completed testing to determine their estimated 1-RM (est. 1-RM) on the barbell back squat and leg extension exercises. Testing procedures conformed to the National Strength and Conditioning Association (NSCA) guidelines for strength testing and were performed by Certified Strength and Conditioning Specialists (CSCS). Testing was performed as follows: i) participants completed a general warm-up of 25 jumping jacks and 10 body weight squats, ii) for a specific warm-up, 50% of participants’ most recent, self-reported back squat max was executed for 10 repetitions, iii) additional warm-up reps were completed at approximately 60% (8-10 repetitions), 70% (5-8 repetitions), 80% (3-5 repetitions), 90% (1-3 repetitions) and 95% (1 repetition) of their self-reported 1-RM. Thereafter, 3-RMs for each exercise were assessed within six attempts, and NSCA guidelines were used to convert the 3-RM loads to est. 1-RM values [27].

#### Hydration testing

Whole-body hydration was assessed through urine specific gravity (USG) analysis. During visits 2 and 4, participants donated a urine sample (~5 mL) that was analyzed using a handheld refractometer (ATAGO; Bellevue, WA, USA). All participants had a USG level ≤ 1.020, and this was used as a threshold of sufficient hydration to continue testing [28].

#### Body composition testing

During visit 2 only, height and body mass were measured using a digital column scale (Seca 769; Hanover, MD, USA). Height was measured to the closest 0.5 cm and body mass to the closest 0.1 kg. Bioelectrical impedance spectroscopy (SFB7; ImpediMed Limited, Queensland, AU) was used to estimate body composition for phenotyping purposes. The procedure was performed according to manufacturer’s instructions. The technique was also explained in more depth by Esco et al. [29] and Moon et al. [30]. After participants rested in a supine position for 5 minutes, two electrodes were placed above and below the right wrist (5 cm apart) and two were placed above and below the anterior portion of the ankle (5 cm apart). Impedance signals were then translated by the device and converted to fat-free mass and fat mass.

#### Vastus lateralis thickness assessments

During visit 2 only, the ultrasound was used to measure VL muscle thickness as reported in Sexton et al. [31] for phenotyping purposes. Participants were instructed to lie supine, with knee and hip fully extended, for a minimum of 5 minutes before image acquisition. Real-time B-mode ultrasonography (NextGen LOGIQe R8, GE Healthcare, Chicago, IL, USA) and a multi-frequency linear-array transducer (L4-12T, 4–12 MHz, GE Healthcare, USA) was utilized to capture a VL image the transverse plane at approximately 50% the distance between the mid-inguinal crease and the proximal patella. The image depth was adjusted until the edge of the femur was in view, which allowed for a full-thickness view of the VL. All ultrasound images were obtained using a generous amount of water-soluble transmission gel. Associated software was used to quantify VL thickness, which was defined as the superficial aponeurosis to the deep aponeurosis. All ultrasound images were taken and analyzed by the same investigator.

#### Skeletal muscle biopsy and processing

During visits 2 and 4, VL biopsies were collected at PRE, 3h POST, and 6h POST using a 5-gauge biopsy needle and local anesthesia. Approximately 50-80 mg of tissue was collected in total. A portion of tissue (~20 mg) was adhered to a cork block via tragacanth gum and preserved in an insulating layer of optimum cutting temperature (OCT) media that was frozen in liquid nitrogen-cooled 2-methylbutane, then stored −80°C to be later used for histology. Remaining tissue was wrapped in prelabelled foils, flash frozen in liquid nitrogen, and stored at −80°C for later use. In total, tissue processing occurred within a 5-minute window.

#### Resistance exercise bouts

During visit 2, participants completed a resistance exercise bout in the morning hours (between 7:00 AM – 10:00 AM) following an overnight fast. The bout format was randomly assigned whereby participants performed 4 sets of the back squat and leg extension exercises at either 80% (80 Fail) or 30% (30 Fail) of their est. 1-RM until volitional failure. At the beginning of the bout participants completed a general warm-up of 25 jumping jacks and 10 body weighted squats, followed by a specific warm-up of 10 repetitions at 30-50% and 5-8 repetitions at 75% of the experimental load assigned to them (i.e., 80% or 30% of est. 1-RM). Participants then initiated sets to failure and were provided 5 minutes of rest between sets. Additionally, participants were given 5 minutes of rest between squats and knee extensions. Failure was determined as either: i) an inability to complete a full repetition, ii) technical errors that potentially compromised safety (e.g., difficulty keeping balance or failure to maintain appropriate posture), or iii) the participant spent longer than 4 seconds between repetitions. Participants reporting nausea, malaise, or light-headedness were provided a cereal bar (calories: 170, total fat: 8 g, carbohydrates: 20 g, protein: 4 g) and a sports drink (calories: 80, carbohydrates: 21 g) to reduce symptoms. After the exercise bout was completed, participants were instructed to return to the laboratory 3 hours and 6 hours later for post-exercise biopsies. Participants were also instructed to not perform rigorous exercise during this time frame.

Seven days later, participants reported back to the laboratory within a two-hour time window of their visit 2 session and performed the bout format that was not performed the week prior. The warm-up, lifting to failure and between-set recovery parameters, and biopsy sampling times were kept identical to the week prior. Additionally, those that were given post-exercise nutrition the week prior were then given the same items following the second exercise bout.

### Biochemical laboratory assays

#### DNA and RNA isolation

Skeletal muscle samples were retrieved from −80°C, crushed on a liquid nitrogen cooled stage, weighed to ~10 mg using a laboratory scale (Mettler-Toledo; Columbus, OH, USA), and added to 500 *μ*L of Trizol in 1.7 mL polypropylene tubes. Muscle was homogenized in Trizol (VWR; Radnor, PA, USA) with tight fitting plastic pestles for approximately 30 seconds and stored in −80°C overnight to increase RNA yield. The following day samples were removed from −80°C and allowed to thaw completely at room temperature. Bromochloropropane (100 *μ*L) was added to samples, samples were shaken for 15 seconds, samples were incubated at room temperature for 2 minutes, and samples were centrifuged at 12,000 g for 15 minutes. Once removed from the centrifuge, samples were kept on ice for the remainder of the procedures. Centrifugation yielded three distinct phases including the top aqueous phase containing RNA, the center meniscus containing DNA, and the bottom phase containing protein. Approximately 200 *μ*L of the aqueous phase was removed and placed in a new tube with 500 *μ*L of 100% isopropanol, and the RNA was subsequently pelleted via centrifugation and reconstituted with DEPC water. Reconstituted RNA was then stored at −80°C and shipped to a commercial laboratory for transcriptome-wide analysis described below.

The remaining DNA and protein were separated by first adding 300 *μ*L of 100% ethanol. Samples were then shaken for 15 seconds, incubated for 3 minutes at room temperature, and subsequently centrifuged at 5,000 g for 10 minutes at room temperature. This produced a DNA pellet and a protein supernatant. The supernatant (protein-Trizol-ethanol mixture) was removed, and the DNA pellet was stored at −80°C until purification was performed using a commercial kit (DNeasy Blood and Tissue Kit, catalog #69504; Qiagen; Germantown, MD, USA) according to manufacturer’s instructions with a minor modification to the elution step. DNA quantity and quality was assessed in duplicate using an absorbance of 260/280 nm provided by a desktop spectrophotometer (NanoDrop Lite; Thermo Fisher Scientific; Waltham, MA, USA). DNA was then stored at −80°C until being shipped on dry ice to a commercial laboratory (TruDiagnostic; Lexington, KY, USA) for the bisulfite conversion and methylation analysis described below.

#### Transcriptome-wide mRNA analysis

RNA was shipped on dry ice to a commercial laboratory (North American Genomics; Decatur, GA, USA) for transcriptomic analysis using the Clariom S Assay Human mRNA array. Raw data were received as .CEL files and analyzed using the Transcriptome Analysis Console v4.0.2 (Thermo Scientific). More details on the statistical analysis are found later in the Methods section.

#### DNA bisulfite conversion and methylome analysis

Bisulfite conversion was performed using an Infinium HD Methylation Assay bisulfite conversion kit (EZ DNA Methylation Kit, Zymo Research, CA, USA). Bisulfite Converted DNA (BCD) was stored at −80°C until DNA methylation analysis. The Infinium MethylationEpic BeadChip Array (Illumina; San Diego, CA, USA) was performed per the manufacturer’s instructions, and BeadChips were imaged using the Illumina iScan^®^ System (Illumina).

#### Nuclear protein isolations

Nuclear protein isolations from frozen muscle tissue were performed using a commercial kit (Nuclear Extraction Kit, catalog# ab113474; Abcam; Waltham, MA, USA). Briefly, skeletal muscle was retrieved from −80°C, crushed on a liquid nitrogen cooled stage, and ~15-20 mg of tissue was added to kit pre-extraction buffer containing DTT. The tissue was then homogenized using tight-fitting pestles, incubated on ice for 15 minutes, and centrifuged for 10 minutes at 12,000 rpm at 4°C. The supernatant was removed, 1x pre-extraction buffer containing DTT and protease inhibitors was added to the pellet, and a 15-minute incubation ensued with 5-second vortexes every 3 minutes. The suspension was centrifuged for 10 minutes at 14,000 rpm at 4°C. The supernatant was transferred to a new tube and protein concentrations were assessed using a bicinchoninic acid (BCA) assay kit (Thermo Fisher Scientific; Waltham, MA, USA) and microplate spectrophotometer (Synergy H1; Biotek; Winooski, VT, USA).

#### Global DNMT and TET activity assays

A DNMT activity assay was performed on nuclear extracts using a commercial assay (Colorimetric DNMT Activity Quantification Kit, catalog# ab113467; Abcam) and microplate spectrophotometer (Synergy H1; Biotek). DNMT activity was expressed as absorbance units per microgram of nuclear protein. The average coefficient of variation for duplicate absorbance values was 4.2%. A TET activity assay was also performed on nuclear extracts using a fluorometric commercial assay (TET Hydroxylase Activity Quantification Kit, catalog# ab156913; Abcam) and microplate fluorometer (Synergy H1; Biotek). TET activity was expressed as fluorescence units per microgram nuclear protein. The average coefficient of variation for duplicate fluorescence values was 7.7%.

#### Immunohistochemistry

Muscle samples that were collected and preserved in OCT as described above were sectioned at a thickness of 10 *μ*m in a cooled (−22°C) cryostat (Leica Biosystems; Buffalo Grove, IL, USA) and electrostatically adhered to positively charged histology slides. The slides were stored at −80°C until immunohistochemical staining whereby were removed from the −80°C, allowed to equilibrate and dry at room temperature for 1.5 hours. Sections were then outlined using a hydrophobic pen to retain solutions for the following incubation steps. First, phosphate buffered saline (PBS) was applied for 10 minutes to rehydrate muscle sections. PBS was then removed, and a 3% peroxide solution was added to sections for 15 minutes. The peroxide solution was removed, and the slides were washed in PBS for three 5-minute washes on a rocker. Sections were then incubated with TrueBlack Lipofuscin Autofluorescence Quencher solution (biotium; Fremont, CA, USA) for 1 minute, and slides were washed in PBS for three 5-minute washes thereafter. The sections were then blocked with a 5% normal goat serum and 2.5% normal horse serum for 1 hour and washed in PBS for 5 minutes. Next, the primary antibody solution was applied (1:20 Mandra, 1:20 of BA_D5, 9:20 and 9:20 5% normal horse serum, all diluted in PBS; all antibodies from Developmental Studies Hybridoma Bank; Iowa City, IA, USA), and slides were incubated overnight in 4°C. The following day, slides were washed four times in PBS (5 minutes per wash) A secondary antibody solution was applied for 1 hour (1:100 goat anti-mouse IgG1 594 and 1:100 goat anti-mouse IgG2B 488 diluted in PBS). Slides were then washed four times in PBS (5 minutes per wash) and a DAPI fluorescent dye (1:10,000 DAPI, diluted in deionized water) was then applied for 15 minutes. Thereafter two 5-minute PBS washes were applied to slides, a 1:1 PBS-glycerol solution was applied around the sections, and glass coverslips were applied thereafter. Immediately following mounting, digital images were captured using a fluorescent microscope (Nikon Instruments, Melville, NY, USA) and a 10x objective. Standardized measurements of type I and type II fiber cross-sectional areas (fCSA) were performed using open-sourced software (MyoVision 2.0) [32,33]. A pixel conversion ratio value of 0.964 µm/pixel was applied to account for the size and bit-depth of images, and a detection range of detection from 500 to 12,000 μm^2^ was used to ensure artifact was removed (i.e., large myofibers which may have not been in transverse orientation, or structures between dystrophin stains which were likely small vessels). Image analysis was also visually inspected to ensure proper analytical fidelity.

### Statistical Analyses

Statistical analyses on training repetition/volume and nuclear activity assays were performed using SPSS v26.0 (IBM Corp, Armonk, NY, USA). Prior to analyses, Shapiro-Wilks tests of normality were performed. Two-way ANOVAs were used to assess main effects and potential interactions of condition (30 Fail vs 80 Fail) x time (PRE, 3h POST, and 6h POST). For any main effect that violated the assumption of sphericity, a Greenhouse-Geiser correction was applied. Significant interactions were further interrogated using dependent samples t-tests between conditions at each time point and within each condition over time.

### DNA methylation analysis

The Partek Genomic Suite V.7 (Partek Inc., MO, USA) was used to process the methylation data as described by Maasar et al. [34]. Briefly, an average detection p-value was assessed for all samples to ensure values below 0.01 [35], and any probes outside of this range were removed from analysis. Raw signals for the methylated and unmethylated probes were assessed for the difference between average median methylated and average median unmethylated probes to ensure the recommended difference of 0.5 or less [35]. After the data was imported to Partek Genomics Suite, single nucleotide polymorphism (SNP) associated probes and cross-reactive probes, identified in validation studies [36], were removed from analysis. Functional normalization using a noob background correction was performed for normalization [37]. Then, principal component analyses (PCA), density plots, and box and whisker plots were used for quality control (i.e., no samples exceeded 2.2 standard deviations or presented abnormal distributions). Following normalization and quality control analysis, differentially methylated position analysis was performed. *β*-values were converted into M-values to represent a more valid distribution of data for analysis of differential methylation [38]. Because both treatments were completed by each participant, paired Samples t-tests were used to assess 30 Fail versus 80 Fail at PRE, 3h POST, and 6h POST, and significant DMPs at PRE were removed as confounding DMPs. An ANOVA was also used to determine the main effects for condition (30 Fail and 80 Fail over time) and for time (PRE vs 3h POST, PRE vs 6h POST, and 3h POST vs 6h POST). Differentially methylated CpG positions (DMPs) were subjected to a significance threshold of p < 0.01. Additionally, an analysis to quantify differentially methylated regions (DMRs) that contained 2 DMPs or more within a short genomic region was performed using the Bioconductor package DMRcate (DOI: 10.18129/B9.bioc.DMRcate). Pathway enrichment analysis of these DMPs was performed using KEGG (Kyoto Encyclopedia of Genes and Genomes) pathways [39–41] via Partek Genomics Suite and Partek Pathway software.

### Transcriptome Analysis

Following the quantification of gene expression with the Clariom S microarray, raw .CEL files were uploaded into the Transcriptome Analysis Console v4.0.2 (TAC) (Thermo Fisher Scientific). The *H. sapiens* genome was used to generate the reference annotations. Two analyses ensued. First, pairwise comparisons were performed to determine mRNAs that were altered from PRE within each bout. A gene target was considered significant if gene expression exceeded a ±1.5-fold-change from PRE and the p-value was less than 0.01. Second, two-way repeated measures (2×2) ANOVAs, with the eBayes correction factor applied, were performed to detect genes that differed between bouts from PRE to 3h POST and PRE to 6h POST. For this statistical analysis, gene expression was considered significant if a fold-change over times between bouts of ±1.5 was exceeded and the interaction p-value was less than 0.01. Bioinformatics using gene lists from both analyses was performed using PANTHER v17.0 [42,43].

### Gene list overlap using KEGG pathways

Gene lists that showed significant DNA methylation and mRNA time effects from PRE to 3h POST, and 6h POST were entered into KEGG pathway analysis [39–41] to examine significantly associated pathways. Thereafter, common pathways that were predicted to be significantly affected from each dataset (p<0.01) were assessed to determine if pathways overlapped.

## RESULTS

### Participant characteristics

Table 1 contains participant characteristics. The male participants that completed the study (n =11) were 23 ± 4 years old with a body mass of 86 ± 12 kg, a height of 180 ± 7 cm, and a BMI of 27 ± 3 kg/m^2^. Resistance training experience (i.e., training age) was 4 ± 3 years and the est. 1RM for the barbell back squat was 143 ± 33 kg (relative to body mass: 1.7 ± 0.3 kg 1RM/ kg body mass). Average vastus lateralis thickness in all participants was 2.99 ± 0.36 cm, and mean fCSA values from biopsied VL tissue averaged 4259 ± 882 *μ*m^2^ (type I fiber percent: 34.6 ± 16.6, type II fiber percent: 65.4 ± 16.6).

**Table 1.**
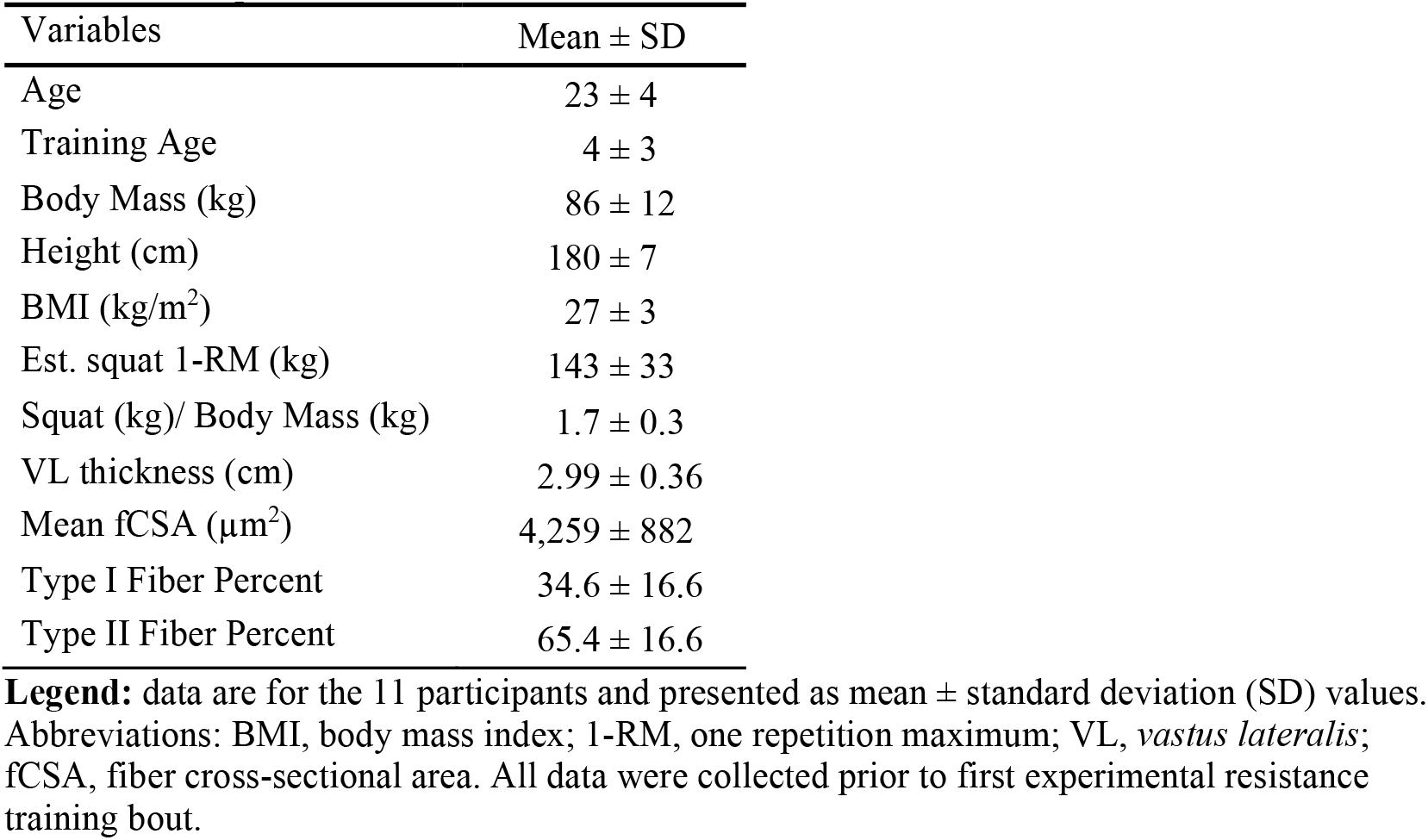
Participant characteristics

| Variables | Mean ± SD |
| --- | --- |
| Age | 23 ± 4 |
| Training Age | 4 ± 3 |
| Body Mass (kg) | 86 ± 12 |
| Height (cm) | 180 ± 7 |
| BMI (kg/m<sup>2</sup>) | 27 ± 3 |
| Est. squat 1-RM (kg) | 143 ± 33 |
| Squat (kg)/ Body Mass (kg) | 1.7 ± 0.3 |
| VL thickness (cm) | 2.99 ± 0.36 |
| Mean fCSA ($\mu\text{m}^2$) | 4,259 ± 882 |
| Type I Fiber Percent | 34.6 ± 16.6 |
| Type II Fiber Percent | 65.4 ± 16.6 |

**Legend:** data are for the 11 participants and presented as mean ± standard deviation (SD) values. Abbreviations: BMI, body mass index; 1-RM, one repetition maximum; VL, *vastus lateralis*; fCSA, fiber cross-sectional area. All data were collected prior to first experimental resistance training bout.

### Training volume differences between the 30 Fail and 80 Fail bouts

Training volume data are presented in Figure 2 (panels A-C). Barbell back squat training volume (Fig. 2A) exhibited a condition by set interaction (p < 0.001) and a main effect of condition (p < 0.001) whereby more volume was performed in the 30 Fail versus 80 Fail condition. A main effect of set was also evident (p < 0.001) whereby average volume decreased across all sets for both conditions. On a per set basis, more back squat volume was performed in the 30 Fail versus the 80 Fail condition (set 1 p < 0.01, set 2 p = 0.002, set 3 p = 0.001, set 4 p = 0.005). Moreover, total back squat volume was greater in the 30 Fail versus 80 Fail condition (p < 0.001). Leg extension training volume exhibited a main effect of condition (p < 0.05, Fig. 2B) where more volume was completed in the 80 Fail versus 30 Fail condition. A dependent samples t-test also indicated that significantly more total leg extension volume was completed in the 80 Fail versus 30 Fail condition (p = 0.030). However, total lower body training volume between bouts was not significantly different (p = 0.570, Fig. 2C).

**Figure 2.**
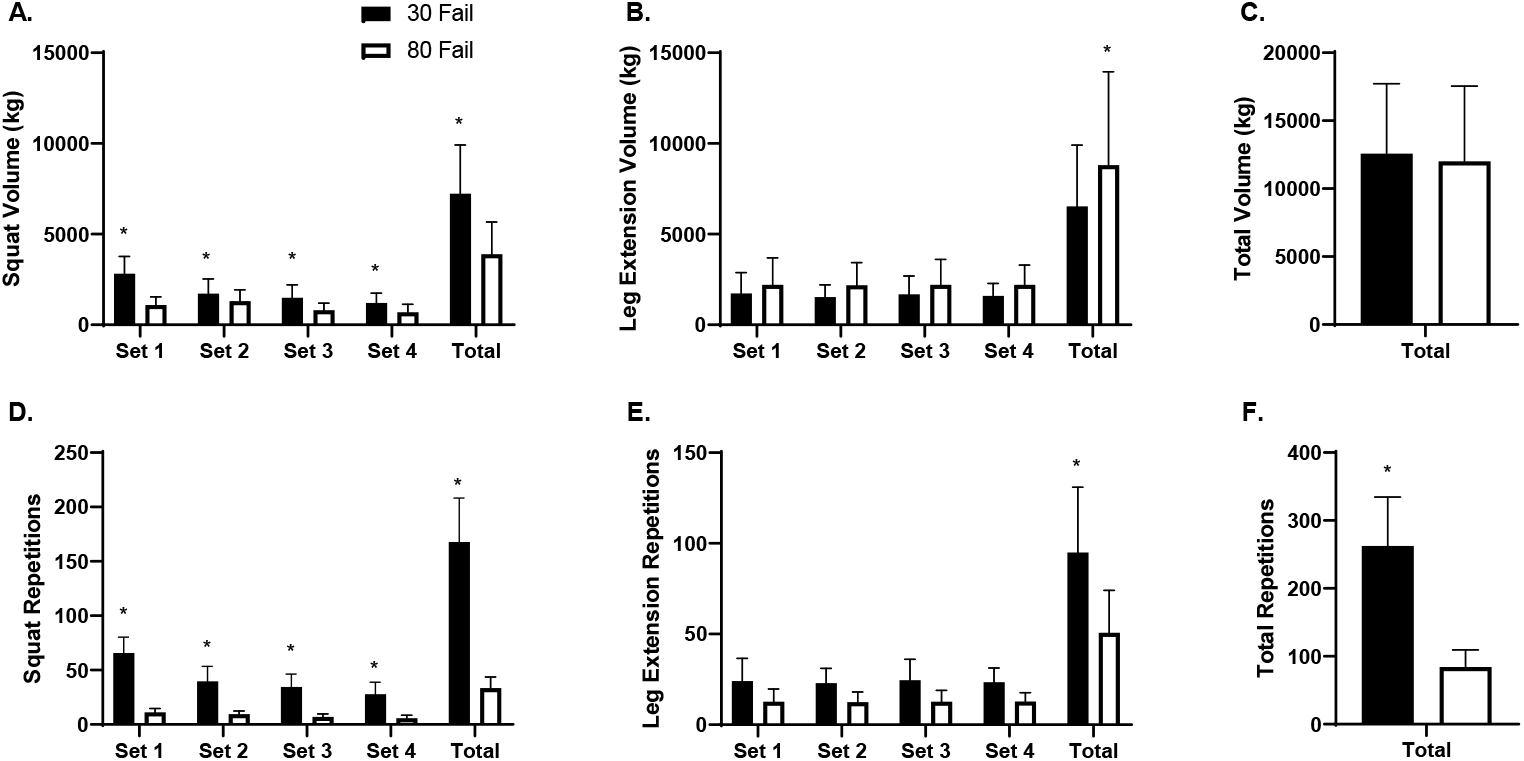
Volume and repetition differences between 30 Fail and 80 Fail bouts **Legend:** Data are presented as mean ± SD for squat volume across sets and total squat volume (panel A), leg extension volume across sets and total leg extension volume (panel B), total volume for both exercises (panel C), squat repetitions across sets and total squat repetitions (panel D), leg extension repetitions across sets and total leg extension repetitions (panel E), and total repetitions performed for both exercises (panel F). Data are for n=11 participants. Symbol: *, indicates significant difference between conditions per set and in totality (p < 0.05). Abbreviations: 30 Fail, 30% one repetition maximum to failure bout; 80 Fail, 80% one repetition maximum to failure bout.

Training repetition data are presented in Figure 2 (panels D-F). The number of back squat repetitions performed exhibited a condition by set interaction (p < 0.001, Fig. 2D) and a main effect of condition (p < 0.001) whereby more repetitions were performed during the 30 Fail versus 80 Fail condition. Additionally, a main effect of set was evident (p < 0.001) whereby repetitions per set decreased in both conditions. Dependent samples t-tests indicated there were significantly more back squat repetitions completed in the 30 Fail versus 80 Fail condition during each set (p < 0.001 for sets 1-4) and for all sets combined (p < 0.001). The repetitions performed across sets during leg extensions showed a main effect of condition (p < 0.001; Fig. 2E) whereby significantly more repetitions were completed per set in the 30 Fail versus 80 Fail condition. Total lower body repetitions completed was also significantly greater in the 30 Fail versus 80 Fail bout (p < 0.001, Fig. 2F).

### DNA methylation changes with the 30 Fail and 80 Fail bouts

Figure 3 presents differentially methylated position (DMP) and region (DMR) data for significant targets (±1.5-fold, p < 0.01), and these data are presented wherein positive values indicate increased methylation in the 30 Fail versus 80 Fail condition and negative values indicate decreased methylation in the 30 Fail versus 80 Fail condition. There were 3,958 significant DMPs between conditions at baseline (p < 0.01). Of these 3,958 DMPs, only 156 possessed a differentially methylated status at 3h POST and/or 6h POST. Therefore, these 156 DMPs were removed from analyses to ensure that changes in each condition were due to the exercise stimuli and not due to altered variation at baseline.

**Figure 3.**
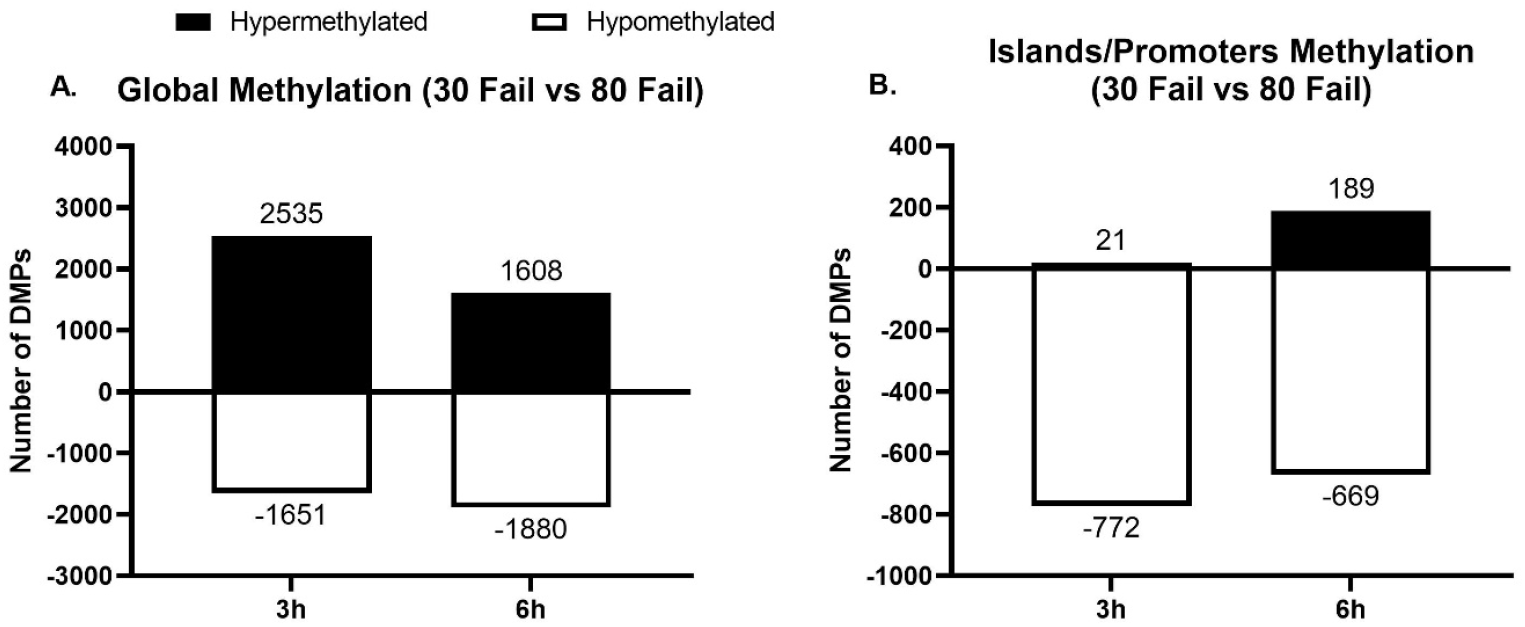
Global and island/promoter methylation differences between bouts **Legend:** Data are presented as total number of significant differentially methylated positions (DMPs) in 30 Fail compared to 80 Fail (panel A), and DMPs present in islands and promoters (panel B). Data are for n=11 participants and significance was established as p < 0.01 between conditions at each time point. Abbreviations: 30 Fail, 30% one repetition maximum to failure bout; 80 Fail, 80% one repetition maximum to failure bout.

At 3h POST there were 4,186 DMPs (first bar in Fig. 3A), with 2,535 DMPs (60.6%) being hypermethylated in 30 Fail versus 80 Fail and 1,651 CpGs (39.4%) being hypomethylated in 30 Fail versus 80 Fail. Of these 4,186 DMPs, 793 DMPs (18.9%) were in a CpG island within a promoter region (first bar in Fig. 3B), and of these 793 DMPs, 21 CpGs (2.6%) were hypermethylated in 30 Fail versus 80 Fail and 772 CpGs (97.4%) were hypomethylated in 30 Fail versus 80 Fail.

At 6h POST there were 3,488 DMPs (second bar in Fig. 3A), with 1,608 DMPs (46.1%) being hypermethylated in 30 Fail versus 80 Fail and 1,880 DMPs (53.9%) being hypomethylated in 30 Fail versus 80 Fail. Of these 3,488 DMPs, 858 DMPs (24.6%) resided in CpG islands within a promoter region (second bar in Fig. 3B), and of these, 189 were hypermethylated in 30 Fail versus 80 Fail (22%) and 669 were hypomethylated in 30 Fail versus 80 Fail (88%).

Regarding DMRs at 3h POST between the 30 Fail and 80 Fail bouts, there were 155 DMRs with 2 or more significant DMPs (p < 0.01), 9 of these DMRs contained 3 DMPs (associated genes: DIP2B, AHCYL, TMEM134, FAM216A, SLC8B1, SERF2, TOM1L2, TEX14, and TXN2), and only 1 DMR contained 4 DMPs (associated gene: EIF4B). Further, 49 of these 155 DMRs (32%) were in CpG islands within promoter regions, with 2 genes containing 3 DMPs (associated genes: FAM216 and TOM1L2) and 1 gene containing 4 DMPs (associated gene: EIF4B).

At 6h POST between the 30 Fail and 80 Fail bouts there were 67 DMRs with at least 2 significant DMPs. Three DMRs contained 3 DMPs (associated genes: SIN3A, RBM39, and PEG10), and only 1 DMR contained 4 DMPs (associated gene: MAP3K3). Of these 67 DMRs, 65 DMRs (97%) were in CpG islands within promoter regions, with these DMRs containing 3 or more DMPs.

### Nuclear TET and DNMT activity

Given robust alterations in DNA methylation with both training bouts, we opted to investigate global TET (demethylating) and DMNT (methylating) activities from nuclear lysates. Due to tissue limitations, n=10 participants yielded enough nuclear lysate material to perform the TET activity assay and n=9 participants yielded enough nuclear lysate material to perform the DNMT activity assay. There was a significant main effect of time for TET activity (p = 0.023, Fig. 4A) where average TET activity increased at both post-exercise timepoints. However, there was no significant main effect of condition (p = 0.163) or condition by time interaction (p = 0.190). For DNMT activity there were no significant main effects of time (p = 0.271), condition (p = 0.096), or condition by time interaction (p = 0.174; Fig. 4B).

**Figure 4.**
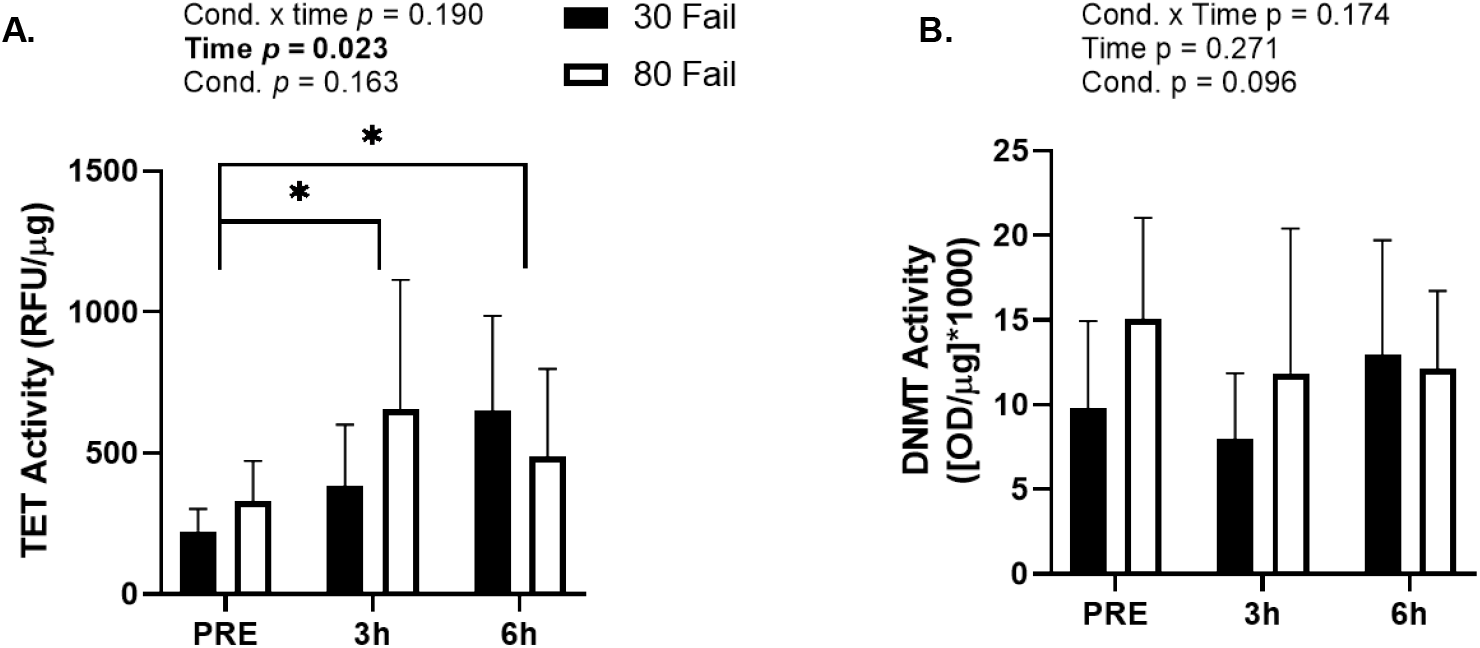
Nuclear TET and DNMT activities between bouts **Legend:** All data are presented as mean±SD for global TET activity (n=10) from PRE to 3h POST and 6h POST resistance exercise (panel A) and DNMT activity (n=9) from PRE to 3h POST and 6h POST resistance exercise (panel B); Symbol: *, indicates a significant change from PRE regardless of bout. Abbreviations: 30 Fail, 30% one repetition maximum to failure bout; 80 Fail, 80% one repetition maximum to failure bout; DNMT, DNA Methyltransferase; OD, optical density; RFU, relative fluorescent units; TET, Ten-Eleven Translocase.

### mRNA expression data and bioinformatics

There were 21,488 genes probed to identify differentially expressed genes (±1.5-fold, p < 0.01; termed ‘DEGs’). There were 889 significant DEGs at PRE between conditions (p < 0.05); thus, these DEGs were removed from analyses to ensure that changes in each condition were due to the exercise stimuli and not due to altered variation at baseline.

From PRE to 3h POST with 30 Fail training 1,428 mRNAs were upregulated and 1,201 mRNAs were downregulated (Fig. 5A). Bioinformatics indicated that mRNAs in the following pathways were significantly enriched (p<0.001, FDR < 0.05): i) Toll receptor signaling (fold-enrichment = 2.78), ii) CCKR signaling (fold-enrichment 2.35), iii) apoptosis signaling (fold-enrichment = 2.31), iv) interleukin signaling (fold-enrichment = 2.19), v) gonadotropin-releasing hormone receptor signaling (fold-enrichment = 1.86), vi) integrin signaling (fold-enrichment = 1.82), vii) inflammation mediated by chemokine and cytokine signaling (fold-enrichment = 1.79). From PRE to 6h POST with 30 Fail training 932 mRNAs were upregulated and 924 mRNAs were downregulated (Fig. 5B). Bioinformatics indicated that mRNAs in the following pathways were significantly enriched: i) CCKR signaling (fold-enrichment 2.44), ii) gonadotropin-releasing hormone receptor signaling (fold-enrichment = 2.13).

**Figure 5.**
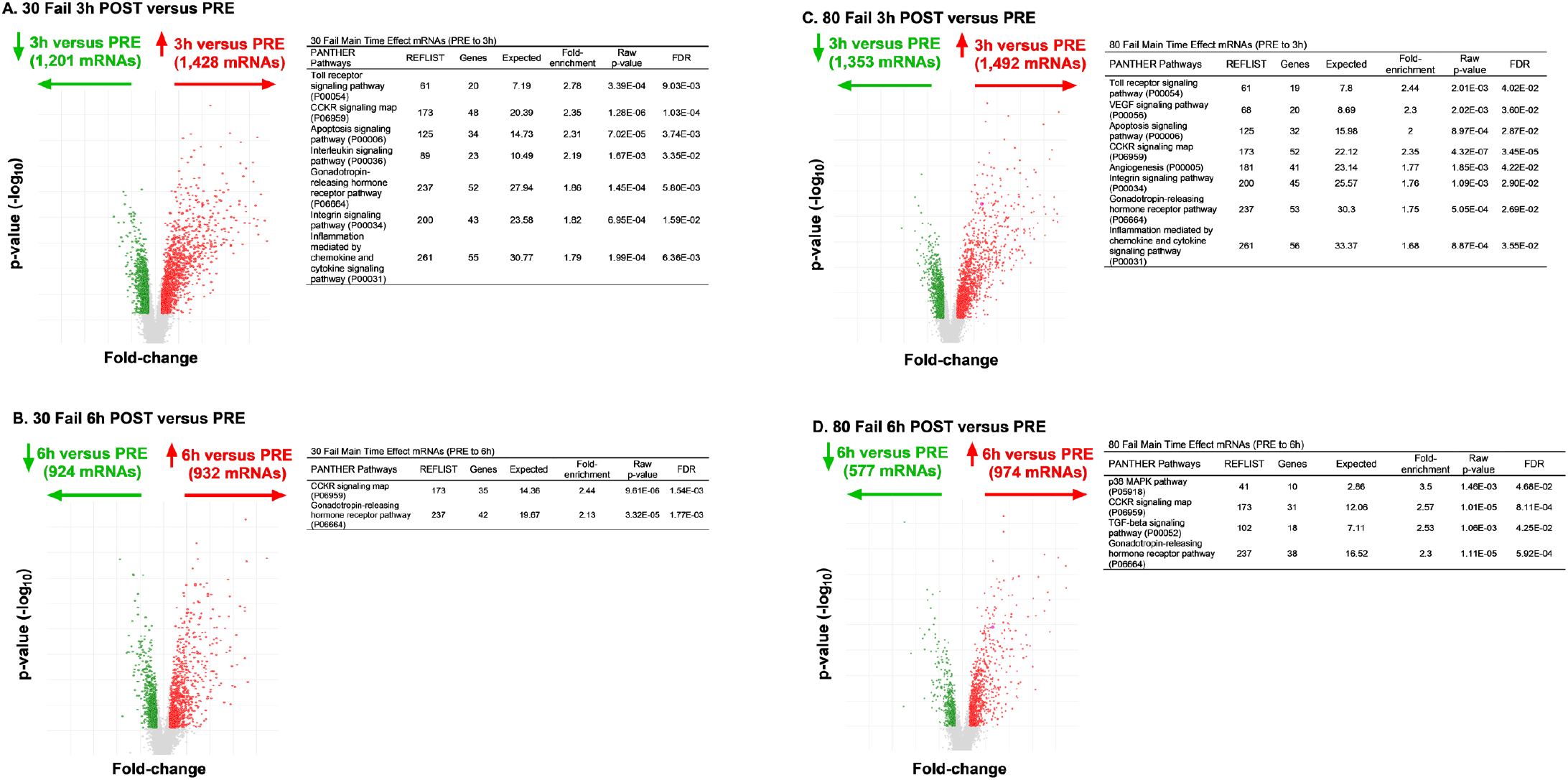
Time effect mRNAs between bouts with pathway enrichment data **Legend:** Differentially expressed genes (DEGs) within the 30 Fail and 80 Fail bouts viewed as volcano plots and PANTHER pathway enrichment. More specifically, 30 Fail training DEGs from PRE to 3h POST (panel A), 30 Fail training DEGs from PRE to 6h POST (panel B), DEGs from PRE to 3h POST with 80 Fail training (panel C), and DEGs from PRE to 3h POST with 80 Fail training (panel D). Data are for n=11 participants, all DEGs have a fold-change from PRE of ±1.5 (p < 0.01), and pathway analysis was significant if False Discovery Rate (FDR) p < 0.05. Abbreviations: 30 Fail, 30% one repetition maximum to failure bout; 80 Fail, 80% one repetition maximum to failure bout; REFLIST, reference gene list across human genome.

From PRE to 3h POST with 80 Fail training 1492 mRNAs were upregulated and 1353 mRNAs were downregulated (Fig. 5C). Bioinformatics indicated that mRNAs in the following pathways were significantly enriched: i) Toll receptor signaling (fold-enrichment = 2.78), ii) VEGF signaling (fold-enrichment = 2.30), iii) apoptosis signaling (fold-enrichment = 2.00), iv) CCKR signaling (fold-enrichment = 2.35), v) angiogenesis (fold-enrichment = 1.77), vi) integrin signaling (fold-enrichment = 1.76), vi) gonadotropin-releasing hormone receptor signaling (fold-enrichment = 1.75), vii) Inflammation mediated by chemokine and cytokine signaling (fold-enrichment = 1.68). From PRE to 6h POST with 80 Fail training 974 mRNAs were upregulated and 577 mRNAs were downregulated (Fig. 5D). Bioinformatics indicated that mRNAs in the following pathways were significantly enriched: i) p38 MAPK signaling (fold-enrichment 2.44), ii) CCKR signaling (fold-enrichment = 2.57), iii) TGF-beta signaling (fold-enrichment = 2.53), iv) gonadotropin-releasing hormone receptor signaling (fold-enrichment = 2.30).

Regarding condition by time interactions (delta-delta fold-change ±1.5, p<0.01), there were only 11 significant DEGs from PRE to 3h POST (Fig. 6A&B), and 17 significant DEGs between bouts from PRE to 6h POST (Fig. 6C&D). Given the low number of differentially expressed genes between bouts over time, no pathways were predicted to differ between bouts at the 3h POST or 6h POST time points.

**Figure 6.**
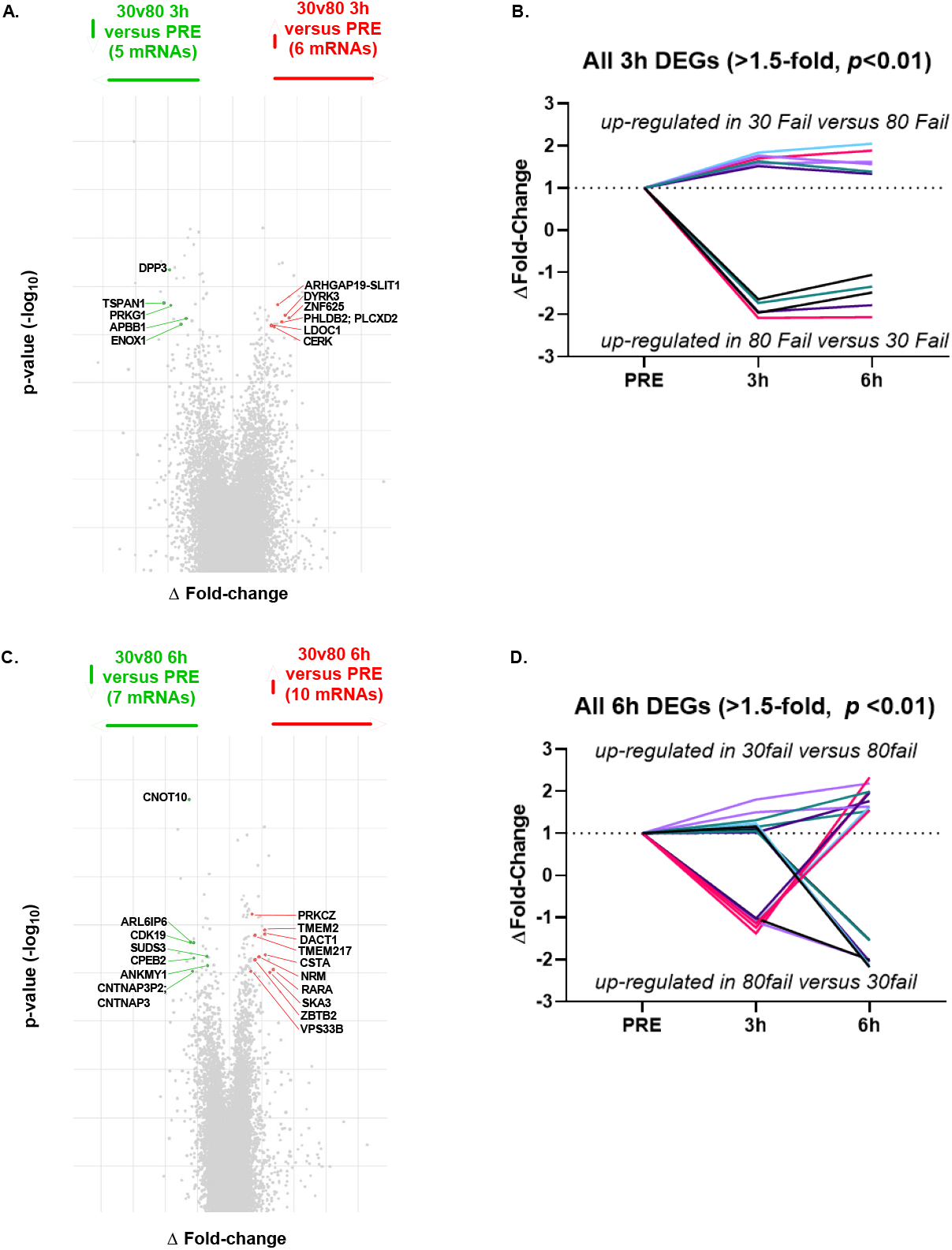
Differentially expressed mRNAs between bouts at both post-exercise time points **Legend:** All data are significant differentially-expressed genes (DEGs) following 30 Fail versus 80 Fail for the ΔFold-change in gene expression plotted against p-value (−log10 transformed) at 3h POST versus PRE (panel A), the ΔFold-change over time for significant DEGs at 3h POST versus PRE (panel B), the ΔFold-change in gene expression plotted against p-value (−log10 transformed) at 6h POST versus PRE (panel C), and the ΔFold-change over time for significant DEGs at 6h POST versus PRE (panel D). Data are for n=11 participants and the significance threshold was established as ±1.5 ΔFold-change, p < 0.01. Abbreviations: 30 Fail, 30% one repetition maximum to failure bout; 80 Fail, 80% one repetition maximum to failure bout.

### Overlapping genome-wide methylome and transcriptome results

An overlay of methylome and transcriptome data can be found in Figure 7. Notably, these analyses only considered mRNAs and DNA methylation values that showed significant main effects of time from PRE.

**Figure 7.**
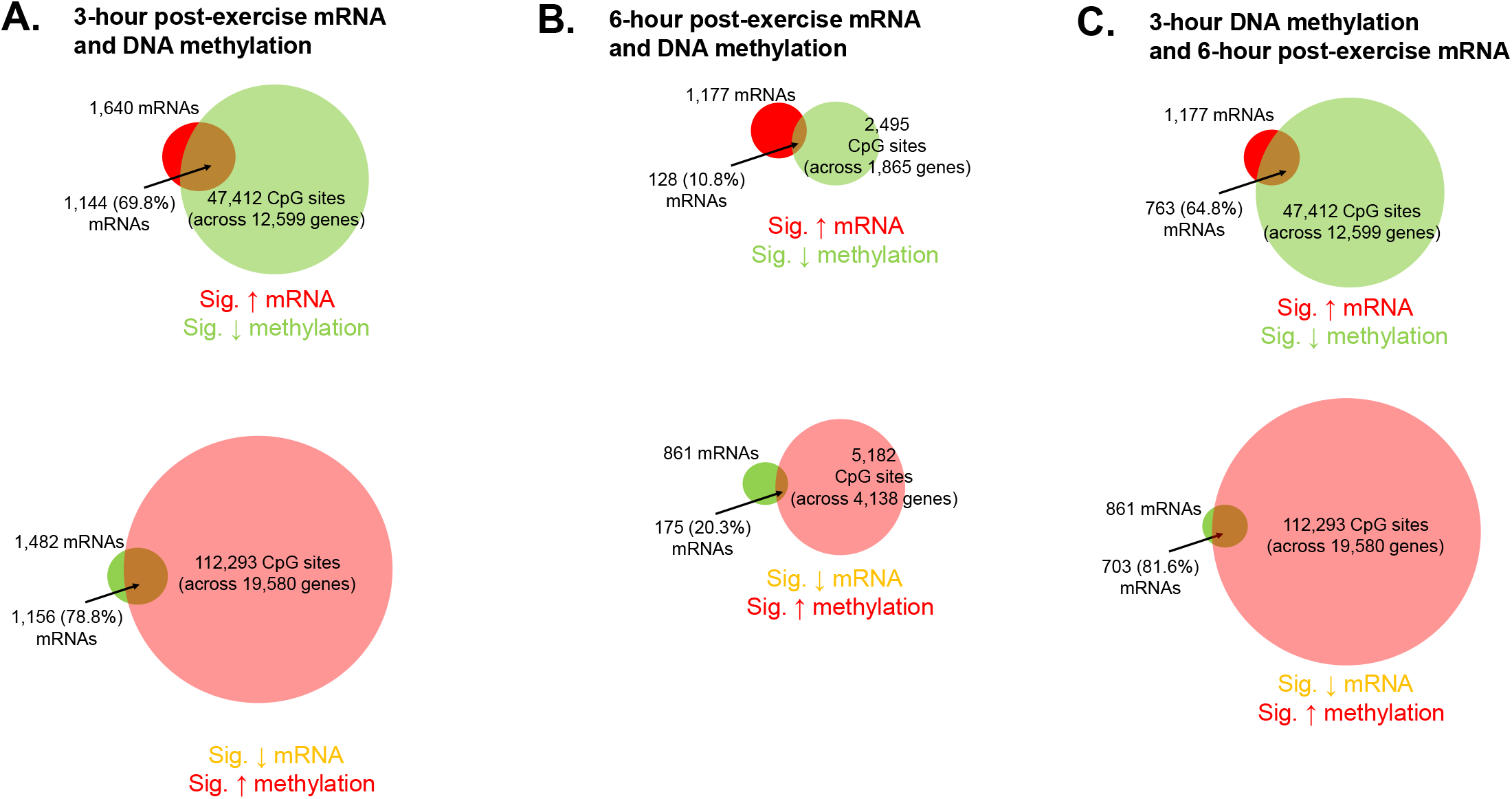
Transcriptome and methylation signatures overlayed **Legend:** Venn diagrams illustrating mRNAs significantly altered 3 hours following exercise (panel A) and 6 hours following exercise (panel B) and their overlap with associated CpG sites that showed inverse methylation patterns. Data are from n=11 participants, differentially methylated CpG sites were considered significantly altered from PRE if p < 0.01, and mRNAs were considered significantly altered from PRE if ±1.5 fold-change (p < 0.01).

A relatively high level of DNA methylation events occurred from PRE to 3h POST, as there was a significant decrease in the methylation status of 69,696 CpG sites and significant increase in 170,255 CpG sites (Fig. 7a). Of the 1,640 mRNAs that were significantly up-regulated at 3h POST, 1,144 (or 68.8%) of these mRNAs had 1 or more associated CpG site that was hypomethylated at this time point. Of the 1,482 mRNAs that were significantly down-regulated 3h POST, 1,156 (or 78.8%) of these mRNAs had 1 or more associated CpG site that was hypermethylated at this time point.

Compared to 3h POST there were fewer DNA methylation events that occurred 6h POST, as there was a significant decrease in the methylation status of 3,578 CpG sites and significant increase in 8,841 CpG sites (Fig. 7b). Of the 1,177 mRNAs that were significantly up-regulated 6h POST, 128 (or 10.8%) of these mRNAs had 1 or more associated CpG site that was hypomethylated at this time point. Of the 861 mRNAs that were significantly down-regulated 6h POST, 175 (or 20.3%) of these mRNAs had 1 or more associated CpG site that was hypermethylated at this time point.

Because DNA methylation precedes alterations in gene transcription, we also investigated DNA methylation changes at 3h POST relative to 6h POST mRNA expression changes (Fig. 7c). Of the 1,177 mRNAs that were significantly up-regulated at 6h POST, 763 (or 64.8%) of these mRNAs had 1 or more associated CpG site that was hypomethylated at 3h POST. Of the 861 mRNAs that were significantly down-regulated 6h POST, 703 (or 81.6%) of these mRNAs had 1 or more associated CpG site that was hypermethylated at 3h POST.

Finally, Table 2 contains KEGG analysis results of significantly altered pathways predicted to be affected following exercise, regardless of bout, according both the DNA methylation and mRNA expression signatures. When removing disease-related pathways from the 3h POST analysis (e.g., “Yersina infection”, “Salmonella infection”, “Cushing syndrome”, etc.), 50 KEGG pathways were significantly enriched (p<0.01) according to the DNA methylation data, 22 KEGG pathways were significantly enriched (p<0.01) according to the mRNA expression data, and 14 (64%) of these pathways overlapped. When removing disease-related pathways from the 6h POST analysis, 40 KEGG pathways were significantly enriched (p<0.01) according to the DNA methylation data, 16 KEGG pathways were significantly enriched (p<0.01) according to the mRNA expression data, and 4 (25%) of these pathways overlapped. Of the 50 KEGG pathways were significantly enriched (p<0.01) according to the 3h POST DNA methylation data and 16 KEGG pathways were significantly enriched (p<0.01) according to the 6h POST mRNA data, 13 (81%) of these pathways overlapped.

**Table 2.**
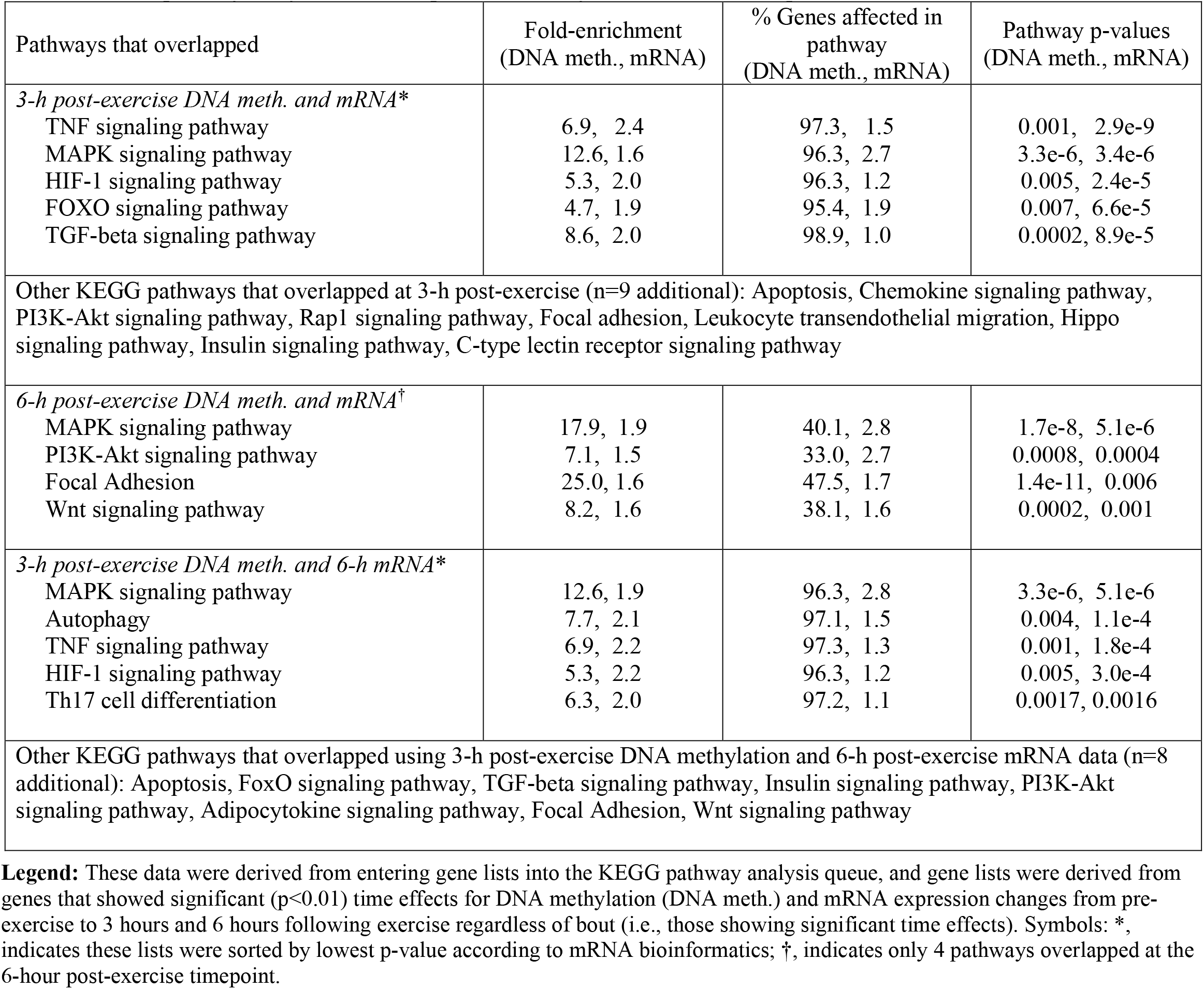
KEGG pathway analysis and overlap of DNA methylation and mRNA expression data

| Pathways that overlapped | Fold-enrichment<br>(DNA meth., mRNA) | % Genes affected in<br>pathway<br>(DNA meth., mRNA) | Pathway p-values<br>(DNA meth., mRNA) |
| --- | --- | --- | --- |
| *3-h post-exercise DNA meth. and mRNA\** |  |  |  |
| TNF signaling pathway | 6.9, 2.4 | 97.3, 1.5 | 0.001, 2.9e-9 |
| MAPK signaling pathway | 12.6, 1.6 | 96.3, 2.7 | 3.3e-6, 3.4e-6 |
| HIF-1 signaling pathway | 5.3, 2.0 | 96.3, 1.2 | 0.005, 2.4e-5 |
| FOXO signaling pathway | 4.7, 1.9 | 95.4, 1.9 | 0.007, 6.6e-5 |
| TGF-beta signaling pathway | 8.6, 2.0 | 98.9, 1.0 | 0.0002, 8.9e-5 |
| Other KEGG pathways that overlapped at 3-h post-exercise (n=9 additional): Apoptosis, Chemokine signaling pathway, PI3K-Akt signaling pathway, Rap1 signaling pathway, Focal adhesion, Leukocyte transendothelial migration, Hippo signaling pathway, Insulin signaling pathway, C-type lectin receptor signaling pathway |  |  |  |
| *6-h post-exercise DNA meth. and mRNA<sup>†</sup>* |  |  |  |
| MAPK signaling pathway | 17.9, 1.9 | 40.1, 2.8 | 1.7e-8, 5.1e-6 |
| PI3K-Akt signaling pathway | 7.1, 1.5 | 33.0, 2.7 | 0.0008, 0.0004 |
| Focal Adhesion | 25.0, 1.6 | 47.5, 1.7 | 1.4e-11, 0.006 |
| Wnt signaling pathway | 8.2, 1.6 | 38.1, 1.6 | 0.0002, 0.001 |
| *3-h post-exercise DNA meth. and 6-h mRNA\** |  |  |  |
| MAPK signaling pathway | 12.6, 1.9 | 96.3, 2.8 | 3.3e-6, 5.1e-6 |
| Autophagy | 7.7, 2.1 | 97.1, 1.5 | 0.004, 1.1e-4 |
| TNF signaling pathway | 6.9, 2.2 | 97.3, 1.3 | 0.001, 1.8e-4 |
| HIF-1 signaling pathway | 5.3, 2.2 | 96.3, 1.2 | 0.005, 3.0e-4 |
| Th17 cell differentiation | 6.3, 2.0 | 97.2, 1.1 | 0.0017, 0.0016 |
| Other KEGG pathways that overlapped using 3-h post-exercise DNA methylation and 6-h post-exercise mRNA data (n=8 additional): Apoptosis, FoxO signaling pathway, TGF-beta signaling pathway, Insulin signaling pathway, PI3K-Akt signaling pathway, Adipocytokine signaling pathway, Focal Adhesion, Wnt signaling pathway |  |  |  |

**Legend:** These data were derived from entering gene lists into the KEGG pathway analysis queue, and gene lists were derived from genes that showed significant ( $p < 0.01$ ) time effects for DNA methylation (DNA meth.) and mRNA expression changes from pre-exercise to 3 hours and 6 hours following exercise regardless of bout (i.e., those showing significant time effects). Symbols: \*, indicates these lists were sorted by lowest p-value according to mRNA bioinformatics; †, indicates only 4 pathways overlapped at the 6-hour post-exercise timepoint.

While not highlighted in Table 2, it is notable that three pathways were predicted to be significantly altered according to the DNA methylation and mRNA signatures in all 3- and 6-hour post exercise comparisons included “MAPK signaling”, “PI3K-Akt signaling”, and “Focal adhesion”.

## DISCUSSION

This study is the first to compare the methylome and transcriptome responses to higher load (80 Fail) versus lower load (30 Fail) resistance exercise bouts. A greater hypomethylation response of CpG sites in promoter regions was observed in the 30 Fail versus 80 Fail condition at both time post-exercise points (3h POST: 772 versus 21, 6h POST: 669 versus 189). Nuclear TET activity significantly increased following exercise regardless of 30 Fail or 80 Fail training. Contrary to our initial hypothesis, transcriptomic data indicated that there were only 11 mRNAs from PRE to 3h POST and 17 mRNAs from PRE to 6h POST that showed significant interactions between bouts. However, there were thousands of mRNAs that were altered following the 30 Fail (Pre to 3h: 2,629 DEGs, Pre to 6h: 1,856 DEGs) and 80 Fail bouts (Pre to 3h: 2,845, PRE to 6h: 1,551), and these gene signatures largely resembled each other according to bioinformatics. Also notable, more robust DNA methylation events occurred 3 hours versus 6 hours following exercise and regardless of bout. Finally, the percentage of significantly altered mRNAs that also demonstrated significantly inversed DNA methylation patterns across one or more CpG sites was greater at 3 hours versus 6 hours following exercise. Finally, because the molecular mechanism of DNA methylation precedes gene transcription high percentages of genes that were up- or downregulated at the later 6-hour post-exercise timepoint also demonstrated significantly inversed DNA methylation patterns across one or more CpG sites at the earlier 3-hour post-exercise timepoint. These findings are discussed in greater detail in this section.

Research interest into how exercise affects genome-wide skeletal muscle DNA methylation has increased in recent years, and multiple associated reviews have been published [25,44–46]. Barres et al. [19] were the first to report that the hypomethylation of various metabolic genes occurs in skeletal muscle following a single high-intensity aerobic exercise session in humans, and that these events corresponded with an alteration in the mRNA expression levels of these genes. A 2021 study from Maasar and colleagues compared the skeletal muscle DNA methylome responses to two bouts of running exercise [34]. The authors reported that CpG sites associated with several metabolic genes exhibited more robust demethylation responses to the higher intensity bout of change-of-direction running versus straight line running. Telles et al. [47] more recently examined how resistance exercise, high-intensity interval exercise, or the combination of both affected the mRNA expression and DNA methylation of select myogenic regulatory factors (MYOD1, MYF5, and MYF6). All exercise protocols were reported to promote DNA demethylation of these genes 4- and 8-hours post-exercise, and the mRNA expression of MYOD1 and MYF6 were elevated 4 hours following exercise. Finally, acute resistance exercise in untrained males has been demonstrated to promote DNA hypomethylation and increased expression of mRNAs associated with matrix/actin structure and remodelling, mechano-transduction, TGF-beta signaling and protein synthesis [22,24].

While these investigations have been insightful, it is difficult to compare our findings to data from these aforementioned studies given the differences in exercise modes and muscle biopsy sampling time points. Notwithstanding, the current study adds meaningful insight to this body of literature. First, the lower load (30 Fail) resistance exercise bout promoted a more robust CpG island/promoter hypomethylation response at both post-exercise time points relative to higher load (80 Fail) resistance exercise. Whether these acute responses eventually translate to differential training adaptations remains to be determined. However, in agreement with the data from Massar and colleagues, this finding supports that implementing different training styles elicits unique post-exercise DNA methylation signatures. Second, a majority of DNA methylation events occurred 3- versus 6-hours post-exercise regardless of bout (239,951 versus 12,419 CpG sites significantly hyper- or hypomethylated at these respective time points, p < 0.01). Indeed, long-lived alterations in skeletal muscle DNA methylation have been reported to occur in response to weeks of resistance training [23], and this phenomenon has been posited to serve as an “epigenetic memory” mechanism to promote more streamlined gene expression responses to subsequent training bouts. However, the data herein also illustrate that a bout of resistance exercise can lead to appreciably rapid and robust genome-wide alterations in skeletal muscle DNA methylation patterns.

Another novel finding resulting from our nuclear lysate assays is the observed changes in nuclear TET, but not DNMT, activity. Comparing our data to other research findings is not possible since this is the first investigation to determine how nuclear DNMT and TET activities are altered by exercise. Indeed, it is plausible that the exercise-induced increase in TET activity observed herein promoted a reduction in DNA methylation. However, there are limitations to this interpretation. First, we were unable to determine which TET isoforms were operative in affecting global TET activity, and it is also possible that the activities of specific DNMT isoforms were differentially affected to result in a nullified post-exercise global DNMT activity change. Second, data presented in Figure 7 indicate that appreciably more CpG sites showed increased methylation relative to CpG sites showing decreased methylation, which implicates that the activities of one or multiple DNMTs was likely elevated during or within the first 3 hours of exercise. Given the novelty and elusiveness of these findings, more refined analytical approaches are needed to determine how exercise affects the activities of various TET and DNMT isoforms. As an aside, there are interesting data linking TET activity to muscle function. In an elegant series of experiments by Wang et al. [48], the authors reported that TET2 knockout mice experienced severe muscle dysfunction. The authors used these data as well as *in vitro* experiments to conclude that TET2 activity is essential for muscle regeneration as well as myoblast differentiation and fusion. Although the implications of these preclinical data are difficult to relate to the current study, the data from Wang and colleagues reiterate the need to examine nuclear TET activity in human skeletal muscle during periods of exercise training and various disease states.

The mRNA response to a single exercise bout does not provide information on other molecular events that promote skeletal muscle anabolism (e.g., translational signaling or the protein synthetic response). Notwithstanding, researchers commonly use bioinformatics to model transcriptomic data and unveil signaling events or pathways that are potentially altered with resistance training [49–51]. Although 30 Fail bout led to a more robust hypomethylation profile of gene promoters as discussed two paragraphs above, the global mRNA expression responses were largely similar between bouts. Specifically, several of the mRNA changes were similar between the 30 Fail and 80 Fail bouts, as only 11 mRNAs differed between conditions from PRE to 3h POST and 17 mRNAs differed between bouts from PRE to 6h POST. Our bioinformatics results also indicated that both training bouts significantly altered mRNAs involved in inflammatory signaling (e.g., Toll receptor signaling, CCKR signaling, chemokine and cytokine signaling), apoptosis signaling, gonadotropin-releasing hormone signaling, and integrin signaling. In extrapolating these bioinformatics data to potential chronic training outcomes, we surmise that the overall phenotypic responses may be largely similar between the 30 Fail and 80 Fail training bouts. There were, however, some differences between bouts worthy of discussion. First, PRKCZ mRNA was up-regulated at 6h POST in the 30 Fail versus the 80 Fail bout, and CDK19 mRNA was up-regulated at 6h POST in the 80 Fail versus the 30 Fail bout. PRKCZ has been shown to bind to and inhibit Akt *in vitro* [52], and Akt acts as a protein kinase to activate the mammalian target of rapamycin (mTOR) signaling cascade to increase muscle protein synthesis [53]. CDK19 is involved with the mediator complex and therefore, along with other proteins, interacts with RNA polymerase II [54]. To our knowledge, neither of these genes have been investigated in skeletal muscle in response to exercise. Hence, it is unknown if the differential post-exercise responses in these two mRNAs between 30 Fail and 80 Fail training are meaningful and/or would lead to different phenotype outcomes. Our bioinformatics analyses also indicated that the VEGF/angiogenesis, p38 MAPK, TGF-beta signaling pathways were predicted to be more affected following the 80 Fail bout, and the interleukin signaling pathway was predicted to be more affected following the 30 Fail bout. VEGF signaling is associated with the formation of new capillaries [55]. Due to the extended time duration needed to complete sets during 30 Fail training, it is counterintuitive that 80 Fail training would preferentially affect these pathways. However, these bioinformatics data remarkably agree with the findings of Campos et al. [56] who reported that participants partaking in eight weeks of resistance training using intermediate repetition ranges (~10/set) experienced significant increase type IIa myofiber capillary number, whereas a separate group utilizing high repetition (~30/set) ranges did not experience this adaptation. The interleukin signaling pathway being significantly enriched following the 30 Fail bout suggests a potential increase in inflammation and myokine formation [57]. It stands to reason 30 Fail training could induce an increased inflammatory response from the greater number of muscle contractions and time under tension [58]. The TGF-beta signaling pathway has been associated with extracellular matrix remodeling in skeletal muscle [59], and the 80 Fail condition perhaps upregulated this gene network due to greater mechanical tension transferred through the extracellular matrix during muscle contraction. Nevertheless, the associations stated above are speculatory given that they are predictions provided by bioinformatics, and further research is needed to determine if these acutely enriched pathways would translate into meaningful adaptations with longer term training paradigms.

When comparing −omics data results in Figure 7, several interesting themes emerge. First, the percentage of significantly altered mRNAs that also demonstrated significantly inversed methylation patterns across one or more CpG sites was appreciably higher at 3 hours versus 6 hours following exercise (~75% versus ~15%, respectively). Moreover, high percentages (65% and 82% respectively) of genes that were up- or downregulated 6 hours following exercise also demonstrated significantly inversed DNA methylation patterns across one or more CpG sites 3 hours following exercise. Only a handful of studies have sought to compare DNA methylation events to corresponding mRNA expression patterns following one or multiple bouts of resistance exercise. For instance, Laker and colleagues [60] reported that only 2% of differentially expressed genes from a one-week (three-bout) resistance training intervention were also differentially methylated. However, these data were confounded by participants being assigned to a high fat diet and the post-intervention biopsy was collected one day following the last exercise bout. Seaborne and colleagues reported that an acute bout of resistance exercise in eight untrained males led to ~10,000 CpG sites becoming hypomethylated and ~7,500 becoming hypermethylated [23]. Interestingly, DNA methylation signatures were not associated with changes in mRNA expression until participants had undergone a chronic training paradigm [23]. Additionally, when acute and chronic methylome and transcriptome data were overlapped, ~40% of differentially expressed genes were shown to be associated with altered DNA methylation signatures [24]. Telles et al. [47] more recently reported that an interrelated, but not time-aligned response, of myogenic regulatory factor gene demethylation and mRNA expression occurred up to 8 hours following resistance exercise and other exercise modalities. However, their analyses were only limited to three genes, and it is notable that all these prior studies examined untrained participants. Notwithstanding, when considering our data in lieu of this prior evidence, it is becoming increasingly evident that resistance exercise-induced alterations in skeletal muscle DNA methylation and mRNA expression patterns are interrelated, albeit this relationship appears to be more coupled rapidly following exercise (i.e., within a 3-hour window) rather than 6+ hours or days following an exercise bout.

Finally, the overlay of DNA methylation and mRNA expression bioinformatics yielded insightful information. As stated in the results, the three pathways were predicted to be significantly altered whereby genes showed overlap at the DNA methylation and mRNA expression levels 3- and 6-hours following exercise included “MAPK signaling”, “PI3K-Akt signaling”, and “Focal adhesion”. Indeed, several studies examining the acute signaling responses to a bout of resistance exercise have implicated that proteins encoded from genes of these pathways show altered phosphoprotein statuses [61–65]. However, data regarding how resistance exercise affects the DNA methylation status and mRNA responses of genes associated with these pathways are sparse. Seaborne and colleagues utilized KEGG pathway analysis with acute and chronic resistance training to report that the skeletal muscle DNA methylation statuses of genes related to “MAPK signaling” and “Focal adhesion” are significantly altered [22]. Turner et al. [24] have overlapped methylome and pooled transcriptome data to report that chronic resistance training affects genes associated with “matrix/actin structure and remodelling”, “Focal adhesion”, and “TGF-beta signalling and protein synthesis”. Hence, the current data continue to provide evidence that post-exercise alterations in DNA methylation and mRNA expression patterns are predicted to affect select genes associated with these cellular processes. Also notable, the pathways predicted to be affected according to 3-hour post-exercise DNA methylation changes and 6-hour post-exercise mRNA expression changes showed the greatest overlap (81%) compared to 3-hour overlay data (64%) and 6-hour overlay data (25%). As has been discussed above, this finding continues to support that earlier DNA methylation events better aligns with mRNA expression events at later time points.

### Experimental Limitations

The present study possesses limitations including a small sample size, the younger healthy male sample population, and muscle sample collection time points. A larger number of more diverse participants and/or muscle samples being collected at additional timepoints across a 24-72h time frame may have yielded more insight.

### Conclusions

In conclusion, 30 Fail and 80 Fail resistance exercise bouts produced unique DNA methylation responses across various gene promoters, albeit largely similar transcriptomic responses. These data continue to add insightful information to the body of literature comparing the muscle-molecular responses of higher load versus lower load resistance training paradigms.

## Acknowledgements

The authors would like to thank the participants for devoting their time and willingness to engage in this study.

